# Mitochondrial membrane tension governs fission

**DOI:** 10.1101/255356

**Authors:** Dora Mahecic, Lina Carlini, Tatjana Kleele, Adai Colom, Antoine Goujon, Stefan Matile, Aurélien Roux, Suliana Manley

## Abstract

During mitochondrial fission, key molecular and cellular factors assemble on the outer mitochondrial membrane, where they coordinate to generate constriction. Constriction sites can eventually divide, or reverse upon disassembly of the machinery. However, a role for membrane tension in mitochondrial fission, although speculated, has remained undefined. We captured the dynamics of constricting mitochondria in mammalian cells using live-cell structured illumination microscopy (SIM). By analyzing the diameters of tubules that emerge from mitochondria and implementing a fluorescence lifetime-based mitochondrial membrane tension sensor, we discovered that mitochondria are indeed under tension. Under perturbations that reduce mitochondrial tension, constrictions initiate at the same rate, but are less likely to divide. We propose a model based on our estimates of mitochondrial membrane tension and bending energy in living cells which accounts for the observed probability distribution for mitochondrial constrictions to divide.

## Introduction

Mitochondria are highly dynamic organelles, transported through the cytoplasm along cytoskeletal networks while they change in size and shape (Nunnari et al., 1997; Youle and van der Bliek, 2012). Mitochondrial dynamics and network connectivity have been linked to bioenergetic function, allowing adaptation of cellular energy production in response to stress (Gomes et al., 2011; Rambold et al., 2011; Tondera et al., 2009) and regulation of the cell cycle (Mitra et al., 2009). Unique among organelles, mitochondria cannot be generated de novo, but instead must proliferate by fission, also referred to as division (Youle and van der Bliek, 2012). Thus, fission plays an important role in ensuring cellular inheritance of mitochondria, as well as being implicated in quality control by acting as a step in the mitophagic pathway (Burman et al., 2017; Twig et al., 2008).

In mammalian cells, the known mitochondrial fission machinery assembles on the outer surface of the organelle. Initially, the fission site is marked by a pre-constriction defined by contact with ER tubules (Friedman et al., 2011) and deformed by targeted actin polymerization (Ji et al., 2015; Korobova et al., 2013; Manor et al., 2015). Subsequently, surface receptors including MiD49/51 (Palmer et al., 2011), Mff (Gandre-Babbe and van der Bliek, 2008; Otera et al., 2010) or Fis1 (Mozdy et al., 2000) accumulate at the preconstriction and recruit dynamin related protein (Drp1) (Labrousse et al., 1999; Smirnova et al., 2001). Drp1 oligomerizes into helices that wrap around the division site, and hydrolyzes GTP to provide a mechano-chemical force for constriction (Fröhlich et al., 2013; Ingerman et al., 2005; Mears et al., 2011; Kalia et al, 2018). In addition, the dynamin 2 protein (Dyn2) can play a role in fission downstream of Drp1 (Lee et al., 2016), albeit a non-essential one (Fonseca et al., 2019; Kamerkar et al., 2018). Interestingly, deformations induced by exogenous mechanical forces can also trigger recruitment of the downstream machinery for mitochondrial fission (Helle et al., 2017). This underlines that membrane fission processes are fundamentally mechanical in nature, triggered by forces that generate membrane deformation.

An additional factor, membrane tension, is known to play a crucial role in other processes involving membrane deformations such as exocytosis (Gauthier et al., 2011), endocytosis (Morlot et al., 2012; Riggi et al., 2019; Roux et al., 2006), cytokinesis (Lafaurie-Janvore et al., 2013), and cell protrusion (Raucher and Sheetz, 2000). Tension in the plasma membrane is a consequence of built-in bilayer tension and stresses from the cytoskeleton and its motors (Keren et al., 2008; Kozlov and Mogilner, 2007), which can together impact the ability of an applied force to drive membrane fission. Indeed, in budding yeast, membrane tethers anchoring mitochondria to the cell cortex were shown to play a role in mitochondrial fission, which was hypothesized to be linked to tension (Klecker et al., 2013). However, the role and origins of membrane tension remain little explored in the context of mitochondrial fission in mammalian cells. This is in part because it is challenging to quantify the tension, even in relative terms, of mitochondria in living cells.

Here, we report a key role for membrane tension in governing mitochondrial division in mammalian cells. Using time-lapse super-resolution imaging, we measured dynamic changes in membrane shape to identify highly constricted sites with diameters below 200 nm. We observed that the presence of the fission machinery, while necessary, is not sufficient to ensure division. We found that constrictions were more likely to result in division when mitochondria were under higher membrane tension. A novel Fluorescence Lifetime Imaging (FLIM) mitochondrial membrane tension sensor (Goujon et al., 2019) revealed that mitochondrial membrane tension was reduced following depolymerization of the microtubule network, a condition that resulted in the same frequency of constriction initiation, but a lower frequency of fissions. Finally, based on our measurements in living cells, we propose a physical model for mitochondrial division in which membrane tension combines with elastic energy during constriction to govern the kinetics and probability of fission.

## Results

### Constriction by the division machinery does not ensure fission

We performed live-cell SIM imaging of COS-7 cells transiently transfected with Drp1-mCherry and GFP targeted to the matrix by the mitochondrial targeting sequence from subunit VIII of human cytochrome c oxidase (mito-GFP). We observed that some constrictions marked by Drp1-mCherry proceeded to fission (Figure 1A, S1, Movie S1), but others lost enrichment of Drp1 without dividing and relaxed to an unconstricted state (termed ‘reversal’) (Figure. 1B, S1, Movie S2). Similar “reversible” or “non-productive” Drp1 constrictions were previously reported in yeast (Legesse-Miller et al., 2003) and mammalian cells (Ji et al., 2015), although their physical properties remained unquantified and their cause was unclear. For quantification purposes, we defined ‘reversals’ as Drp1-enriched constriction sites that reached a diameter below 200 nm before relaxing, well below the mean mitochondrial diameter of ~500 nm (Figure 1C,D, S1).

**Figure 1:**
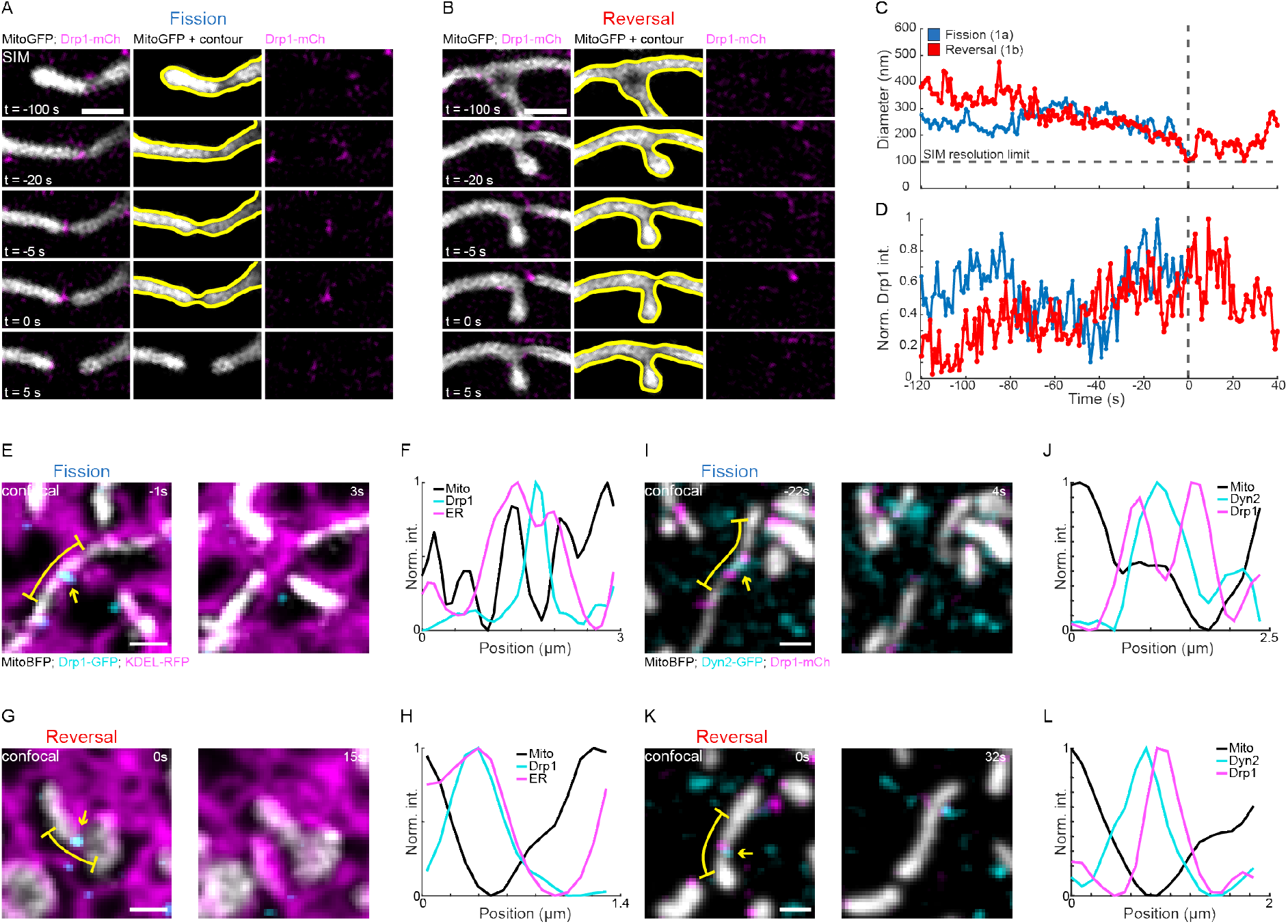
Key molecular components are present at constriction sites that undergo reversals. (A, B) Time-lapse SIM imaging of COS-7 cells transiently transfected with mito-GFP (greyscale) and mCherry-Drp1 (magenta) showing an example (A) fission and (B) reversal (see Movies S1,2, SFig 1). (C) Time evolution of diameter at the mitochondrial constriction site measured for fission (blue) and reversal (red) events shown in A, B. (D)

Integrated intensity of Drp1 at the constriction site over time measured for fission (blue) and reversal (red) events shown in A, B, normalized to the maximum value. (E, G) Time-lapse live-cell confocal imaging of Mito-BFP, Drp1-GFP and KDEL-RFP showing examples of fission and reversal. (F, H) Intensity profiles of ER, Mito and Drp1 intensities adjacent to the yellow dashed lines in E,G. (I, K) Time-lapse SIM imaging of mitochondria and Dyn2 showing a reversal at a Dyn2-enriched constriction site. (J, L) Intensity profiles of ER, Mito and Drp1 intensities adjacent to the yellow dashed lines in I, K. Scale bar represents 500 nm in A and B. Scale bar represents 1 μm in E, G, I, K, and yellow arrows mark the constriction site.

To examine whether reversals result from differences in fission machinery, we imaged several fission factors – the ER, Drp1 and Dyn2, restricting our analysis to Drp1-mediated constrictions. Since our SIM imaging was limited to two colors, we performed fast (1 Hz), three-color live-cell confocal imaging of mitochondria and Drp1, with either Dyn2 or the ER (Figure 1E-L). We found that Dyn2 could be present or absent at Drp1-mediated constrictions (Figure 1I-L), with 30% of fissions and 36% of reversals enriched in Dyn2 (N=30 and 33 respectively) (Figure 1I-L). These observations are consistent with recent reports that Drp1, but not Dyn2 is essential for mitochondrial division (Kamerkar et al., 2018). We further measured colocalization between Drp1 mediated mitochondrial constrictions and ER tubules, as such contacts were shown to mark sites prior to division (Friedman et al., 2011). We found that both fissions and reversals could occur at constrictions overlapping with ER tubules (90% for fissions (N=10) and 89% for reversals (N=18), Figure 1E,G), which appear as peaks in intensity profiles along the constriction site (Figure 1F,H).

An accumulation of Drp1 at these sites typically coincided with an increased rate of constriction, measured at ~17 nm/s for fissions and 18 nm/s for reversals during the 5 seconds leading up to maximal constriction, suggesting active constriction by Drp1 (Figure S2, N=61 for fissions and N=38 for reversals). Some sites underwent several cycles of constriction and relaxation, coupled with Drp1 accumulation and disassembly (Figure 1D). Cyclic dynamics could lead to either fission or reversal, 3±2 constriction cycles/min (N=61) and 2±1 cycles/min respectively (N=38), implying that neither abundance of Drp1, nor constriction rate, nor cyclic activity distinguishes fissions from reversals. Overall, 66% of constriction sites underwent fission (N=112, Figure S1), while the remaining 34% ended as reversals (N=57, Figure S1). Thus, differences in observed constriction dynamics or machinery do not account for differences in success to divide.

### Fission events are characterized by increased membrane tension

The division machinery wraps around mitochondria, providing a force that locally constricts the organelle. However, in other examples of membrane fission, an interplay between force and membrane tension determines whether fission is driven to completion (Gauthier et al., 2011; Lafaurie-Janvore et al., 2013; Morlot et al., 2012; Raucher and Sheetz, 2000; Riggi et al., 2019; Sinha et al., 2011). We noticed that after division, daughter mitochondria would recoil away from the division site (Figure 2A), reminiscent of an elastic body being cut under tension (Movie S4, S5). Therefore, we decided to examine the relationship between membrane tension and the probability of fission versus reversal.

**Figure 2:**
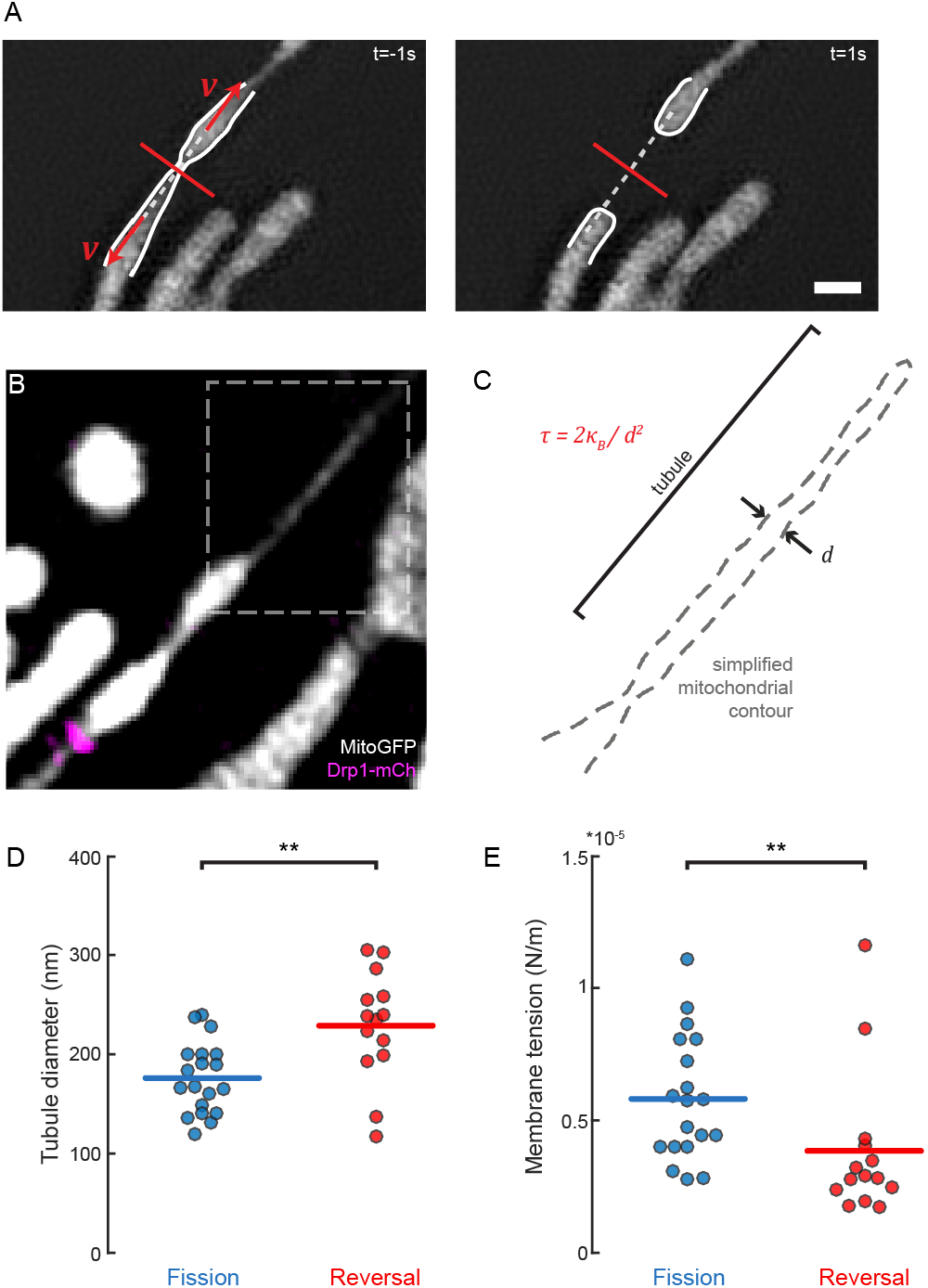
Estimated membrane tension for fissions and reversals. (A) Time-lapse SIM images of a mitochondrion (mito-GFP) 1 sec before (left) and after (right) fission showing the recoil of daughter mitochondria post fission. Measured retraction velocities v (red arrows) were projected perpendicular to the constriction site (white dashed line). (B) Fluorescence image showing a constricted mitochondrion with a pulled membrane tube (boxed region). Scale bar: 1 μm. (C) Mitochondrial contour from the outlined region in (A) showing the diameter of the tube d, used as a readout for tension τ. (D) Distribution of tubule radii measured between fission and reversal events. (E) Distribution of calculated membrane tension values between fission and reversal events. Statistical significance calculated by 1- and 2-tailed Mann-Whitney U test where appropriate: **P<0.01.

In vitro experiments can estimate membrane tension by pulling on a membrane and measuring the size of the resulting membrane tubule (Derényi et al., 2002; Evans and Yeung, 1994). Analogously, microtubule motors in living cells can spontaneously extrude mitochondrial membrane nanotubes (Huang et al., 2013; Wang et al., 2015) (Figure 2B). We observed nanotubes just before fissions or reversals, at similar frequencies (19% and 24% for N=101 and 59 respectively). We inferred the membrane tension by classical energy minimization, which gives a relationship between the nanotube diameter, membrane tension *τ*, and membrane bending rigidity *K_B_* (Derényi et al., 2002; Evans and Yeung, 1994) (Figure 2C, SI):

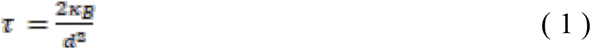

The average diameters of tubules pulled from mitochondria that subsequently either divided or reversed were 176 ± 4 nm and 229 ± 7 nm respectively (Figure 2D). Thus, the population of mitochondria undergoing fission was on average under significantly higher membrane tension at 5.81 ± 0.54 x 10^-6^ N/m, compared to the population undergoing reversals at 3.85 ± 0.75 x 10^-6^ N/m (Figure 4e, N = 19 and N = 14 respectively, mean ± SEM). We found consistent values when analyzing the recoil motion of mitochondria post-fission to independently estimate membrane tension (SI, Figure S3). Together, these data show that mitochondrial constrictions which are under higher tension are more likely to undergo fission.

**Figure 3:**
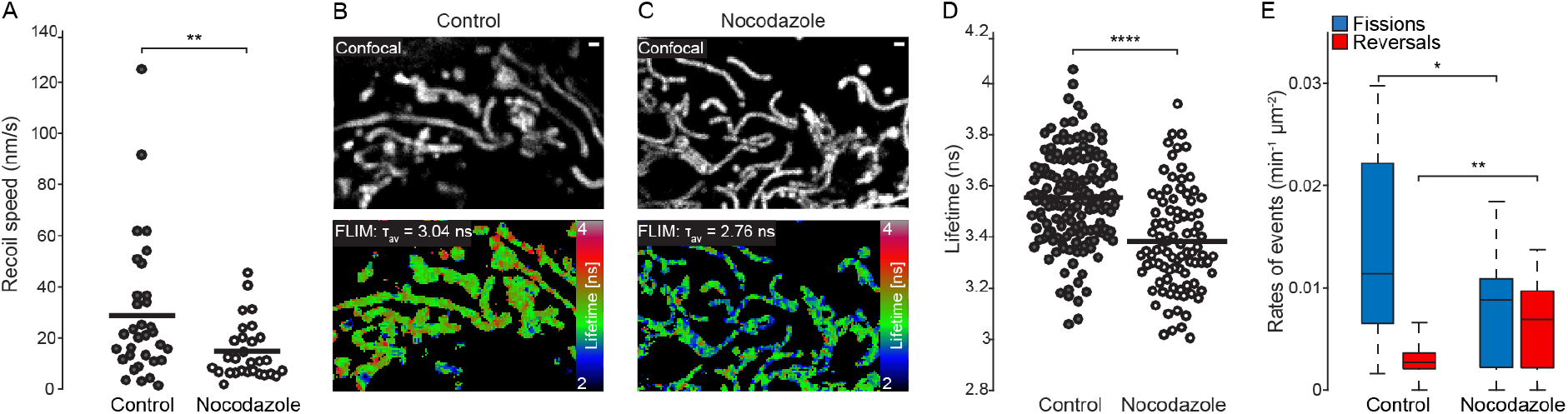
Membrane tension under nocodazole perturbation. (A) Distribution of recoil speeds of mitochondria post-division, between control and nocodazole treated cells. (B,C) Confocal (top) and FLIM (bottom) images of mitochondria in cells stained with the mitochondria-targeted FliptR fluorescent tension probe under (B) control conditions and (C) nocodazole treatment. (D) Distributions of bulk fluorescence lifetimes of the FliptR fluorescent tension probe between control and nocodazole-treated cells. (E) Box chart showing rates of fission and reversal in Nocodazole treated (N=16) and control (N=16) cells. Scale bars: 1μm. Statistical significance calculated by 1- and 2-tailed Mann-Whitney U test where appropriate: *P<0.05, **P<0.01, ****P<0.0001.

**Figure 4:**
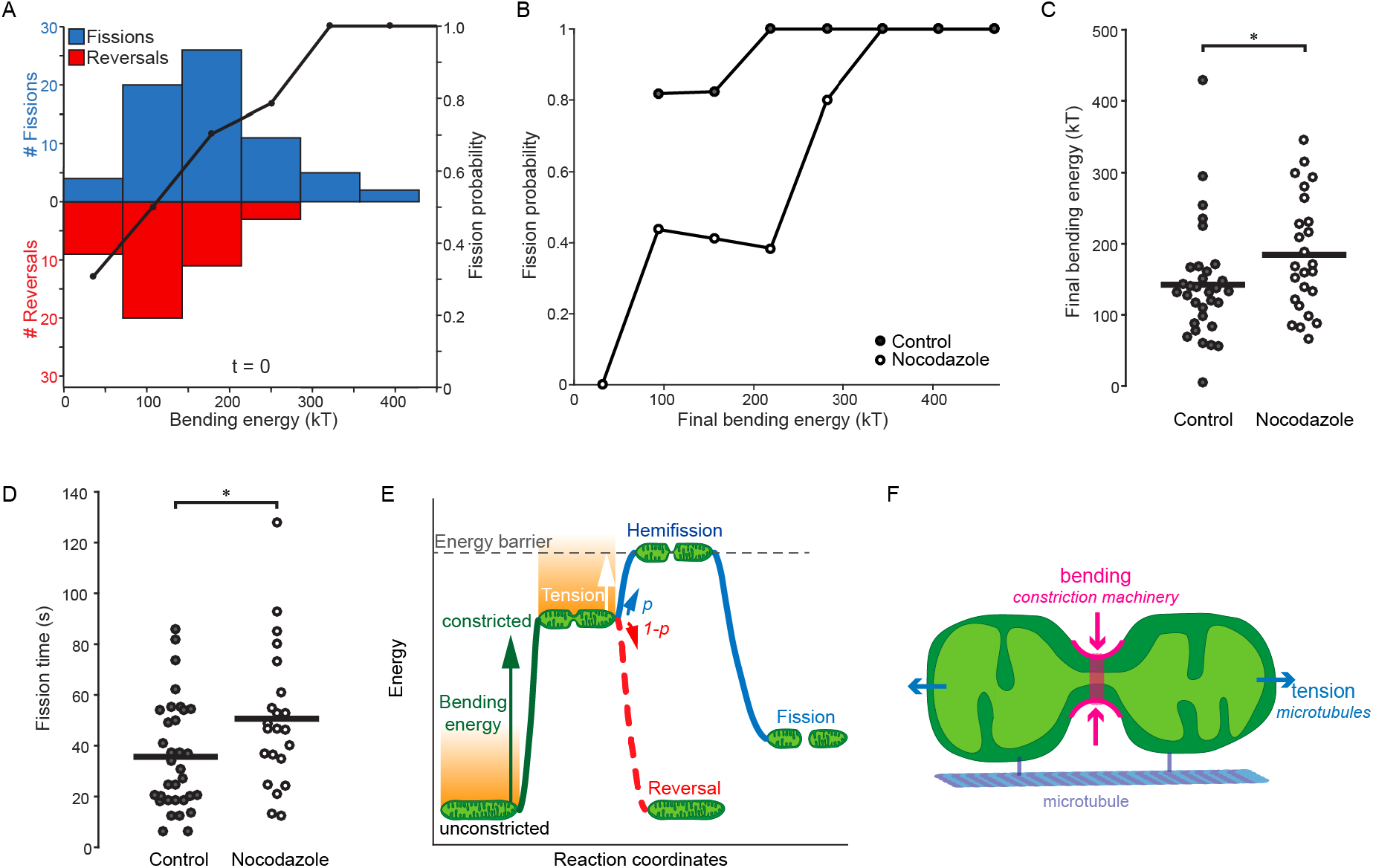
Fission timing and probability related to bending energy and tension. (A) Left: Histogram showing numbers of fissions and reversals at different local bending energy intervals. Right: Experimental probability of fission calculated as ratio of fissions to total constrictions at different local bending energy intervals. (B) Experimental probability of fission of control and nocodazole treated cells. (C) Distribution of final bending energies between control and nocodazole treated fission events. (D) Distribution of fission times between control and nocodazole treated fission events. (E) Cartoon of the probabilistic model of mitochondrial fission showing the contribution of bending energy (green line) and membrane tension (orange shaded area) in reaching the energy barrier for fission (grey dashed line). Both bending energy and tension set the probability p of fission (blue line). Reversals occur either due to a lack of bending energy or low probability of necessary fluctuation energies. (F) Schematic representation of the different contributions to fission probability: bending energy (magenta) and tension (blue). Statistical significance calculated by 1- and 2-tailed Mann-Whitney U test where appropriate: *P<0.05.

Additionally, membrane tubes under tension can spontaneously develop undulations through a “pearling instability”. We observed undulations on 11% of dividing mitochondria (N=88) (Figure S3), also previously reported during mitochondrial fragmentation (Gonzalez-Rodriguez et al., 2015) and in neuronal mitochondria (Cho et al., 2017). We found that constricted mitochondria exhibiting pearling modes eventually underwent fission of at least one of the constriction sites (100%, N=10). Conversely, reversals of pearling modes were rarely observed (4%, N=57) and occurred exclusively following fission at a neighboring constriction site, suggesting that the loss of tension released during fission could be responsible (Figure S3, Movie S4, S5).

### Reduced membrane tension results in increased probability of reversals

We set out to understand the origin of membrane tension in dividing mitochondria. Post-division recoil suggests the presence of external forces pulling or anchoring mitochondria. Mitochondrial transport is mainly mediated through microtubule motors (Boldogh and Pon, 2007), and we hypothesized that they could generate membrane tension. To test this hypothesis, we depolymerized microtubules using nocodazole (De Brabander et al., 1976; Hoebeke et al., 1976). Indeed, during a timeframe of 1-hour post-treatment while cell and organelle morphologies were maintained (Figure S3), we observed a decrease in recoil velocities (Figure 3A) consistent with a reduction of estimated membrane tensions by 40% (SI, N=33 control and N=26 nocodazole).

To more directly test whether depolymerizing microtubules decreases membrane tension, we used a mitochondrial-targeted variant of the mechanosensitive FliptR probe (Figure 3B,C, Figure S3) (Colom et al., 2018; Goujon et al., 2019; Soleimanpour et al., 2016). The fluorescence lifetime of FliptR depends on the orientation between its chromophoric groups, which is sensitive to membrane tension. Comparing control versus nocodazole-treated cells, FliptR showed significantly shorter average fluorescence lifetimes, indicative of an overall reduction in mitochondrial membrane tension (Figure 3D).

Having established that nocodazole treatment reduces mitochondrial membrane tension, we examined its consequences on mitochondrial division by quantifying the probability for constriction sites to divide or reverse. Importantly, the rate of Drp1-induced constrictions initiated per mitochondrial area was unperturbed by nocodazole treatment (~ 0.014 min^-1^ μm^-2^). Furthermore, the degree of overlap between mitochondria and ER remained unchanged (Figure S3), as did mitochondrial diameter and membrane potential (Figure S3), suggesting that the mitochondrial physiology and the ability of the division machinery to constrict were unaffected by nocodazole treatment. However, we found a 2.4-fold increase in the rate of reversal events, and a concomitant decrease in the rate of fission (Figure 3E). Thus, a reduced membrane tension did not change the initiation of Drp1-mediation constrictions, but reduced the efficiency of their fission significantly.

### Tension modifies the energy landscape of mitochondrial fission

The fission process can be represented as an energy landscape where during fission, the elastic energy stored in the mitochondrial membrane overcomes an energy barrier *E_f_*. During constriction, the local elastic energy increases through tension and bending of the membrane. To account for membrane bending and its contribution to the elastic energy of the membrane, we estimated the membrane bending energy of mitochondrial constrictions from their shape (Figure S4, SI) and by numerically evaluating the Helfrich equation (Helfrich, 1973). Both fissions and reversals accumulated bending energy at the constriction, reaching 188±14 k_B_T versus 127±10 k_B_T respectively at maximal constriction (mean±SEM, N=70 and 43 respectively, Figure S4, Appendix A). Note that our ability to estimate the bending energy is limited by the resolution of the contour. We find that there is significant overlap between the distributions (Figure S4), and a range of values of constricted state bending energies can result in either outcome (Figure S4), underlining the probabilistic nature of this model.

To estimate the energy barrier to fission, we calculated the probability of fission *p*(*E*) defined experimentally as the ratio of the number of fissions to all constrictions with a given energy. *p*(*E*) increases with local bending energy, and by determining the bending energy at which all constrictions result in fission (*p*(*E*)=1), we could estimate the energy barrier to fission as ~300 k_B_T Considering that mitochondria are double-membraned organelles, this estimate is consistent with simulations of dynamin-mediated scission (Morlot et al., 2012), as well as theoretical estimates for a hemifission state, which spontaneously leads to fission (Kozlovsky and Kozlov, 2003).

To test the effect of membrane tension on the energy landscape for fission, we compared the experimental probability of fission between control and nocodazole treated cells. We found it was shifted towards higher bending energies when membrane tension is reduced (Figure 4B). Therefore, achieving a similar probability of fission would now require more deformation to increase the energy of the constricted state (Figure 4C, N=33 control and 22 nocodazole). We also noticed that Drp1 appeared to reside for longer time periods at mitochondrial constriction sites in nocodazole-treated cells. Fission events in nocodazole-treated cells required on average ~12±7 s longer (Figure 4D, N=33 control and N=22 nocodazole), reflecting decreased fission probability and consistent with a major role for membrane tension in driving the final step of fission.

## Discussion

We report that membrane tension plays a key role in governing mitochondrial fission. How might tension promote mitochondrial division? In a model proposed for dynamin-mediated endocytosis, thermal fluctuations of the membrane bring it over the energy barrier to fission (Morlot et al., 2012). One possibility is that tension fluctuations play an analogous role to thermal fluctuations to overcome the fission barrier. Microtubule-dependent motor proteins, which anchor and transport mitochondria along microtubules, were shown to generate piconewtons of force on millisecond timescales (Carter and Cross, 2005). This suggests that tension can be modulated several orders of magnitude faster than it takes a mitochondrion to divide.

In such a tension-driven model, when the division machinery induces constriction, it brings membranes closer to the energy barrier to fission (Figure 4E,F). Fluctuations in tension could then stochastically deform mitochondrial membranes, storing additional elastic energy at the constriction site to overcome the energy barrier to fission. According to such a model, since constrictions cannot be maintained indefinitely, mitochondrial constriction sites that do not experience a large enough fluctuation during their lifetime will become reversals. This model is supported by our independent estimates of the energy contribution from tension and the magnitude of the mean fluctuation energy extracted from the experimental probability of fission, both of which are ~100 k_B_T (SI). We note that this estimate indicates that thermal fluctuations are insufficient for mitochondrial fission. Furthermore, considering our observations in nocodazole treated cells, our model suggests that lower membrane tension results in lower fluctuation energy (SI), and thus increases the probability of reversal.

Overall, the proposed probabilistic nature of mitochondrial fission may play a role in regulating mitochondrial network morphologies. For instance, mitochondrial division has been observed to take place near replicating nucleoids (Lewis et al., 2016) – the presence of which might create ‘rigid islands’ that alter the mechanical properties at adjacent constriction sites (Feng and Kornmann, 2018), making them more likely to divide according to our model. Such internal mechanisms could simultaneously control the positioning and fate of mitochondrial constrictions. Furthermore, our work suggests how remodeling of the microtubule cytoskeleton could impact global mitochondrial morphology and proliferation through changes in mitochondrial membrane tension. Additional work is needed to examine the role of microtubules in establishing mitochondrial membrane tension and their regulation during mitochondrial fission.

## Acknowledgements

We would like to thank Hélène Perreten for technical assistance, Dr. Hari Shroff for the mitoGFP construct and Dr. Gia Voeltz for the Drp1 and Dyn2 constructs. We would like to thank Niklas Berliner, Jennifer Lippincott Schwartz, Simon Schutz and Dr. Tobias Schneider for helpful discussions. Imaging data used in this publication was produced in collaboration with the Advanced Imaging Center, a facility jointly supported by the Gordon and Betty Moore Foundation and HHMI at HHMI’s Janelia Research Campus. We thank Lin Shao and Teng-Leong Chew at Janelia AIC for their help with SIM imaging. We thank Timo Rey, Sofia Zaganelli, Benoit Kornmann, Qian Feng, Thomas Misgeld and Leanne Godinho for comments on the manuscript. Research in S.M.’s and A.R.’s laboratories is supported by the National Centre of Competence in Research Chemical Biology. T.K. received funding from European Molecular Biology Organization (ALTF-739-2016) and the Munich Cluster for Systems Neurology (SyNergy).

## Author contributions

Conceptualization, D.M., L.C., T.K., A.R., S.M.; Methodology, D.M., L.C., T.K., A.R., S.M.; Software, D.M., L.C.; Validation, D.M., L.C., T.K.; Formal Analysis, D.M., L.C., A.C.; Investigation, D.M., L.C., T.K., A.C.; Resources, A.C., A.G., S.Mat., A.R.; Data Curation, D.M. L.C.; Writing – Original Draft, D.M., L.C., S.M.; Writing – Review and Editing, D.M., L.C., T.K., A.C., A.G., S.Mat., A.R., S.M.; Visualization, D.M., L.C., T.K.; Supervision, A.R., S.M.; Project Administration S.M.; Funding Acquisition, T.K., S.M.

## Materials and Methods

### Cell culture, transfections and dye labelling

Cos-7 cells were grown in Dulbecco’s modified Eagle medium (DMEM) supplemented with 10% fetal bovine serum (FBS). Cells were plated on 25 mm, #1.5 glass coverslips (Menzel) 16-24 h prior to transfection at a confluency of ~10^5 cells per well. Dual transfections containing mCh-Drp1 (Addgene, plasmid #49152) and Mito-GFP (gift from Hari Shroff, Cox8a presequence) were performed with either Lipofectamine 2000 (Life Technologies) or using electroporation (BioRad Xcell). Lipofectamine transfections were carried out in Opti-MEM using 150 ng of mCh-Drp1, 150 ng of Mito-GFP and 1.5 μL of Lipofectamine 2000 per 100 μL Opti-MEM. Electroporation was performed using salmon sperm as a delivery agent. Briefly, cells were pelleted by centrifugation and resuspended in OPTI-MEM. Plasmids and sheared salmon sperm DNA were added to 200 μL of the cell suspension prior to electroporation using a Bio-Rad Gene Pulser (190 Ω and 950 μFD).

Triple transfections containing mCh-Drp1, Mito-BFP (Addgene, plasmid #49151) and Dyn2-GFP (gift from Gia Voeltz) were performed with Lipofectamine 2000. Such transfections were carried out in using 80 ng of mCh-Drp1, 100 ng of Dyn2-GFP and 80 ng of Mito-BFP and 1.5 μL of Lipofectamine 2000. Dual color imaging of dynamin was performed using double transfections of either 100 ng Dyn2-GFP and 150 ng Mito-Scarlet, or 100 ng Dyn2-mCherry and 150 ng Mito-GFP. Triple transfection containing Mito-BFP, Drp1-GFP and KDEL-RFP were performed with Lipofectamine 2000. Such transfections were performed using 100 ng Mito-BFP, 100 ng Drp1-GFP and 100 ng KDEL-RFP. All quantities listed are per well of cells containing 2 mL of culture medium and carried out with Opti-MEM. The Lipofectamine mixture sat for 20 min before its addition to cells.

### Drug treatment

Nocodazole was diluted to a stock solution of 10 mM in DMSO. To depolymerize microtubules, cells were incubated with 10μM Nocodazole (Sigma-Aldrich) for 1h before imaging (1 μL Nocodazole per 1000 μL medium). Control cells were incubated with the equivalent volume of DMSO for 1h before imaging (1 μL DMSO per 1000 μL medium).

### SIM imaging and reconstruction

Fast dual-color SIM imaging was performed at Janelia Farm with an inverted fluorescence microscope (AxioObserver; Zeiss) using an SLM (SXGA-3DM; Fourth Dimension Displays) to create the illumination pattern and liquid crystal cell (SWIFT; Meadowlark) to control the polarization. Fluorescence was collected through a 100X 1.49 NA oil immersion objective and imaged onto a digital CMOS camera (ORCA-Flash4.0 v2 C11440; Hamamatsu). Timelapse images were acquired every 1 s for 3-5 min, with 50 ms exposure time. Fast dual color imaging of mitochondria and Drp1 was performed at 37°C with 5% CO2, in pre-warmed DMEM medium. Dual-color SIM imaging for Nocodazole and Dyn2 experiments was performed on an inverted fluorescence microscope (Eclipse Ti; Nikon) equipped with an electron charge coupled device camera (iXon3 897; Andor Technologies). Fluorescence was collected with through a 100x 1.49 NA oil immersion objective (CFI Apochromat TIRF 100XC Oil; Nikon). Images were captured using NIS elements with SIM (Nikon) resulting in temporal resolution of 1 s for single-color and 6-8s for dual-color imaging, with 50 ms exposure time. Imaging was performed at 37°C in pre-warmed Leibovitz medium. See SI for details on iSIM imaging, image reconstruction and analysis.

SIM images were reconstructed using a custom 2D linear SIM reconstruction software obtained at Janelia farm, as previously described (Gustafsson, 2000; Gustafsson et al., 2008). Images were reconstructed using a generalized Weiner filter parameter value of 0.02-0.05 with background levels of ~100.

### iSIM imaging and reconstruction

For iSIM experiments, imaging was performed on a custom-built microscope setup as previously described (York et al., 2013). The microscope was equipped with a 1.49 NA oil immersion objective (APONXOTIRF; Olympus), with 488 nm and 561 nm excitation lasers and an sCMOS camera (Zyla 4.2; Andor). Images were captured at 0.1-0.3 s temporal resolution for both channels. All imaging was performed at 37°C in pre-warmed Leibovitz medium. Raw iSIM images were deconvolved using the Lucy-Richardson deconvolution algorithm (Lucy, 1974; Richardson, 1972) implemented in MATLAB, run for 40 iterations.

### Confocal imaging

Confocal imaging was performed on an inverted microscope (DMI 6000; Leica) equipped with hybrid photon counting detectors (HyD; Leica). Fluorescence was collected through a 63x 1.40 NA oil immersion objective (HC PL APO 63x/1.40 Oil CS2; Leica). Images were captured using the LAS X software (Leica). All imaging was performed at 37°C in prewarmed Leibovitz medium.

### STORM imaging and reconstruction

For STORM imaging, prior to staining, cells were washed with PBS (Sigma). Cells were incubated with MitoTracker Red CMXRos (LifeTechnologies) at a concentration of 500 nM for 5 minutes, before washing again with PBS.

For measuring mitochondrial membrane potential, cells were incubated with 100 nM TMRE (Abcam, ab113852) for 10 minutes before time-lapse measurements.

STORM imaging was performed at room temperature in a glucose-oxidase/catalase (Glox) oxygen removal buffer described in (Shim et al., 2012). Briefly, a 2% glucose solution is prepared in DMEM (Gibco). Glucose oxidase (0.5 mg/mL) and catalase (40 μg/mL) were added to the glucose solution and the pH was left to drop for 30-60 min. After this time, the pH was adjusted to 7 yielding a final solution with 6.7% HEPES. Imaging was performed on an inverted microscope (IX71; Olympus) equipped with a 100x NA 1.4 oil immersion objective (UPlanSAPO100X; Olympus) using an electron multiplying CCD camera (iXon+; Andor Technologies), with a resulting pixel size of 100nm. Laser intensities were between 1-5 kWcm-2.

For STORM datasets, single molecules were localized using the RapidSTORM v3.3 software (Wolter et al., 2012). Local signal-to-noise detection with a threshold value of 50 was used. Peaks with a width between 70-300 nm and at least 200 photons were rendered for the final STORM image.

### FliptR synthesis

The FliptR probe was synthesized following previously reported procedures (Colom et al., 2018). For mitochondrial targeting, compounds 2,3 and 5 were synthesized and purified according to procedures that will be reported elsewhere in another manuscript (Goujon et al., 2019) in due time (SFigure 7).

Compound 5 was synthesized and purified according to procedures described in (Goujon et al., 2019) (SFigure 7).

The probe can report on membrane tension as reported in reference (Colom et al., 2018). Spectroscopic characterizations, mechanosensitive behavior in LUVs and GUVs of various lipid composition, colocalization studies in mitochondria and response of fluorescence lifetime to osmotic shocks (i.e. membrane tension changes) is reported in (Goujon et al., 2019).

### FLIM imaging and analysis

For FLIM imaging with the mitochondria-targeted FliptR probe, cells were incubated with 500 nM of the probe solution for 15 min, and washed before imaging. Imaging was performed using a Nikon Eclipse TI A1R microscope equipped with a time-correlated single-photon counting module from PicoQuant. A pulsed 485 nm laser (PicoQuant LDH-D-C-485) was used for excitation, operated at 20 MHz. The emission was collected through ha 600/50 nm bandpass filter, on a gated PMA hybrid 40 detector and a PicoHarp 300 board (PicoQuant).

FLIM data was analyzed using the SymPhoTime 64 software (PicoQuant). The fluorescence decay data was fit to a double exponential model after deconvolution for the calculated impulse response function. The values reported in the main text are the average lifetime intensity.

### Statistics

Statistics were performed using Matlab and OriginPro software. All datasets were tested for normal distribution using the D’Agostino-Pearson normality test (significance value of 0.05) (Antonio Trujillo-Ortiz, 2015). If the datasets passed the test, then statistical significance was determined using a two-tailed t-tests. If datasets failed the normality test, a nonparametric test was chosen to compare the significance of means between groups Mann-Whitney test for two samples (with one or two tailed distributions where appropriate) and Kruskal-Wallis ANOVA for multiple samples (Giuseppe Cardillo, 2015). P<0.05 were considered as significant and were marked by ‘*’; P<0.01 with ‘**’, P<0.001 by ‘***’ and P<0.0001 by ‘****’.

Curve fitting was performed using the curve fitting toolbox in Matlab.

