## Supplementary Information for "Mitochondrial membrane tension governs fission"

### Membrane bending energy and tension govern mitochondrial division

**This PDF file includes:**

Supplementary text
Figs. S1 to S6
Captions for movies S1 to S5
References for SI reference citations

**Other supplementary materials for this manuscript include the following:**

Movies S1 to S5

### Materials and Methods

#### Cell culture, transfections and dye labelling.

Cos-7 cells were grown in Dulbecco's modified Eagle medium (DMEM) supplemented with 10% fetal bovine serum (FBS). Cells were plated on 25 mm, #1.5 glass coverslips (Menzel) 16-24 h prior to transfection at a confluency of ~10^5 cells per well. Dual transfections containing mCh-Drp1 (Addgene, plasmid #49152) and Mito-GFP (gift from Hari Shroff, Cox8a presequence) were performed with either Lipofectamine 2000 (Life Technologies) or using electroporation (BioRad Xcell). Lipofectamine transfections were carried out in Opti-MEM using 150 ng of mCh-Drp1, 150 ng of Mito-GFP and 1.5 μL of Lipofectamine 2000 per 100 μL Opti-MEM. Electroporation was performed using salmon sperm as a delivery agent. Briefly, cells were pelleted by centrifugation and resuspended in OPTI-MEM. Plasmids and sheared salmon sperm DNA were added to 200 μL of the cell suspension prior to electroporation using a Bio-Rad Gene Pulser (190 Ω and 950 μFD). Triple transfections containing mCh-Drp1, Mito-BFP (Addgene, plasmid #49151) and Dyn2-GFP (gift from Gia Voeltz) were performed with Lipofectamine 2000. Such transfections were carried out in using 80 ng of mCh-Drp1, 100 ng of Dyn2-GFP and 80 ng of Mito-BFP and 1.5 μL of Lipofectamine 2000. Dual color imaging of dynamin was performed using double transfections of either 100 ng Dyn2-GFP and 150 ng Mito-Scarlet, or 100 ng Dyn2-mCherry and 150 ng Mito-GFP. Triple transfection containing Mito-BFP, Drp1-GFP and KDEL-RFP were performed with Lipofectamine 2000. Such transfections were performed using 100 ng Mito-BFP, 100 ng Drp1-GFP and 100 ng KDEL-RFP. All quantities listed are per well of cells containing 2 mL of culture medium and carried out with Opti-MEM. The Lipofectamine mixture sat for 20 min before its addition to cells.

#### Drug treatment.

Nocodazole was diluted to a stock solution of 10 mM in DMSO. To depolymerize microtubules, cells were incubated with 10μM Nocodazole (Sigma-Aldrich) for 1h before imaging (1 μL Nocodazole per 1000 μL medium). Control cells were incubated with the equivalent volume of DMSO for 1h before imaging (1 μL DMSO per 1000 μL medium).

#### SIM imaging and reconstruction.

Fast dual-color SIM imaging was performed at Janelia Farm with an inverted fluorescence microscope (AxioObserver; Zeiss) using an SLM (SXGA-3DM; Fourth Dimension Displays) to create the illumination pattern and liquid crystal cell (SWIFT; Meadowlark) to control the polarization. Fluorescence was collected through a 100X 1.49 NA oil immersion objective and imaged onto a digital CMOS camera (ORCA-Flash4.0 v2 C11440; Hamamatsu). Time-lapse images were acquired every 1 s for 3-5 min, with 50 ms exposure time. Fast dual color imaging of mitochondria and Drp1 was performed at 37˚C with 5% CO2, in pre-warmed DMEM medium. Dual-color SIM imaging for Nocodazole and Dyn2 experiments was performed on an inverted fluorescence microscope (Eclipse Ti; Nikon) equipped with an electron charge coupled device camera (iXon3 897; Andor Technologies). Fluorescence was collected with through a 100x 1.49 NA oil immersion objective (CFI Apochromat TIRF 100XC Oil; Nikon). Images were captured using NIS elements with SIM (Nikon) resulting in temporal resolution of 1 s for single-color and 6-8s for dual-color imaging, with 50 ms exposure time. Imaging was performed at 37˚C in pre-warmed Leibovitz medium. See SI for details on iSIM imaging, image reconstruction and analysis. SIM images were reconstructed using a custom 2D linear SIM reconstruction software obtained at Janelia farm, as previously described (1, 2). Images were reconstructed using a generalized Weiner filter parameter value of 0.02-0.05 with background levels of ~100.

#### iSIM imaging and reconstruction.

For iSIM experiments, imaging was performed on a custom-built microscope setup as previously described (3, 4). The microscope was equipped with a 1.49 NA oil immersion objective (APONXOTIRF; Olympus), with 488 nm and 561 nm excitation lasers and an sCMOS camera (Zyla 4.2; Andor). Images were captured at 0.1-0.3 s temporal resolution for both channels. All imaging was performed at 37˚C in pre-warmed Leibovitz medium. Raw iSIM images were deconvolved using the Lucy-Richardson deconvolution algorithm (5, 6) implemented in MATLAB, run for 40 iterations.

**Confocal imaging.**
Confocal imaging was performed on an inverted microscope (DMI 6000; Leica) equipped with hybrid photon counting detectors (HyD; Leica). Fluorescence was collected through a 63x 1.40 NA oil immersion objective (HC PL APO 63x/1.40 Oil CS2; Leica). Images were captured using the LAS X software (Leica). All imaging was performed at 37˚C in pre-warmed Leibovitz medium.

**STORM imaging and reconstruction.**
For STORM imaging, prior to staining, cells were washed with PBS (Sigma). Cells were incubated with MitoTracker Red CMXRos (LifeTechnologies) at a concentration of 500 nM for 5 minutes, before washing again with PBS.
For measuring mitochondrial membrane potential, cells were incubated with 100 nM TMRE (Abcam, ab113852) for 10 minutes before time-lapse measurements.
STORM imaging was performed at room temperature in a glucose-oxidase/catalase (Glox) oxygen removal buffer described in Shim et al (7). Briefly, a 2% glucose solution is prepared in DMEM (Gibco). Glucose oxidase (0.5 mg/mL) and catalase (40 µg/mL) were added to the glucose solution and the pH was left to drop for 30-60 min. After this time, the pH was adjusted to 7 yielding a final solution with 6.7% HEPES. Imaging was performed on an inverted microscope (IX71; Olympus) equipped with a 100x NA 1.4 oil immersion objective (UPlanSAPO100X; Olympus) using an electron multiplying CCD camera (iXon+; Andor Technologies), with a resulting pixel size of 100nm. Laser intensities were between 1-5 $kWcm^{-2}$.
For STORM datasets, single molecules were localized using the RapidSTORM v3.3 software (8). Local signal-to-noise detection with a threshold value of 50 was used. Peaks with a width between 70-300 nm and at least 200 photons were rendered for the final STORM image.

**FliptR synthesis.**
The FliptR probe was synthesized following previously reported procedures (9). For mitochondrial targeting, compounds 2,3 and 5 were synthesized and purified according to procedures that will be reported elsewhere in another manuscript {Goujon et al., *manuscript in preparation*} in due time (SFig. 3a).
Compound 5 was synthesized and purified according to procedures described in {Goujon et al., *submitted*} (SFig. 3a).
The probe can report on membrane tension as reported in reference (9). Spectroscopic characterizations, mechanosensitive behavior in LUVs and GUVs of various lipid composition, colocalization studies in mitochondria and response of fluorescence lifetime to osmotic shocks (i.e. membrane tension changes) will be reported elsewhere in another manuscript {Goujon et al., *submitted*} in due time.

**FLIM imaging and analysis.**
For FLIM imaging with the mitochondria-targeted FliptR probe, cells were incubated with 500 nM of the probe solution for 15 min, and washed before imaging. Imaging was performed using a Nikon Eclipse TI A1R microscope equipped with a time-correlated single-photon counting module from PicoQuant. A pulsed 485 nm laser (PicoQuant LDH-D-C-485) was used for excitation, operated at

20 MHz. The emission was collected through ha 600/50 nm bandpass filter, on a gated PMA hybrid 40 detector and a PicoHarp 300 board (PicoQuant).
FLIM data was analyzed using the SymPhoTime 64 software (PicoQuant). The fluorescence decay data was fit to a double exponential model after deconvolution for the calculated impulse response function. The values reported in the main text are the average lifetime intensity.

**Statistics.**
Statistics were performed using Matlab and OriginPro software. All datasets were tested for normal distribution using the D'Agostino-Pearson normality test (significance value of 0.05) (10). If the datasets passed the test, then statistical significance was determined using a two-tailed t-tests. If datasets failed the normality test, a nonparametric test was chosen to compare the significance of means between groups Mann-Whitney test for two samples (with one or two tailed distributions where appropriate) and Kruskal-Wallis ANOVA for multiple samples (11). $P<0.05$ were considered as significant and were marked by '*'; $P<0.01$ with '**', $P<0.001$ by '***' and $P<0.0001$ by '****'.
Curve fitting was performed using the curve fitting toolbox in Matlab.

**Image analysis.**
Images were first segmented using the open-source software ImageJ/Fiji (http://fiji.sc) and the Weka Segmentation V3.2.17 plugin with the resulting probability map used as the segmented image. Subsequent analysis was performed using a custom MATLAB functions which contoured the mitochondria and created a backbone with a mesh (Supplementary Fig. 6). This allowed us to measure the diameter and curvature along the constricted mitochondrion, and hence estimate the local bending energy. Tracking the position of the constriction site allowed us to measure the local Drp1 intensity. Tracking the leading edge of divided daughter mitochondria was used to estimate the membrane tension prior to fission.
The custom MATLAB package can be found on the Github repository (https://github.com/LEB-EPFL/MitoWorks) and allowed us to perform the summarized steps (Supplementary Fig. 6):

1. *mitoTrack*: Identify mitochondria in the segmented image and find nearest neighbor mitochondria in the subsequent frame. Mitochondrial tracks are made up of nearest neighbour mitochondrial centers of mass (using the knnsearch function) in consecutive frames. The tracked mitochondrion of interest is then cropped from both segmented and original time-lapse images for different channels
2. *genContour*: Create contour of the mitochondrion of interest using a built-in Matlab function based on Chan-Vase active contouring of the segmented (12). A small Gaussian filter is applied prior to contouring. The generated contour is then smoothed over a length scale of ~ 170 nm, which was found to be optimal for eliminating noise without sacrificing envelope curvature sensitivity. Contour smoothing was performed using a third-party function based on least-squares smoothing for MATLAB (13).
3. *mitoMesh*: Create a backbone, or centerline, of the mitochondrion of interest. The backbone is smoothed over a lengthscale of ~ 150 nm. Use generated backbone to divide mitochondrion into smaller segments, with the boundaries represented by a mesh, defined by lines drawn perpendicular to the backbone with subpixel spacing (14).
4. *genCurv*: measure the curvature along the contour of the mitochondrion using a third party 'LineCurvature2D' function for MATLAB (15).
5. *minDiameterSearch*: Use the diameters measured along the mesh to track the position of the constriction site, and measure its diameter.
6. *measureBE*: Use mesh to measure dimensions of individual segments. For each segment, use the measured diameters and envelope curvatures to estimate the bending energy and bending energy density of that segment. Find the length scale that maximizes the local bending energy density.

7. *genFWHM*: Generate FWHM based contour by fitting profiles plotted along the mesh with a Gaussian function. Connect the measured widths at along the mitochondrion and smooth at a length scale of ~ 170 nm to generate the FWHM contour.
   Repeat minDiameterSearch and measureBE for FWHM contour.
8. *measureDrp1*: Measure the local Drp1 integrated intensity within a chosen radius around the constriction site. Radius used for all datasets: ~500 nm. Subtract background and bleach correct using a custom-written linear bleach correction function.
9. *mitoPull*: Repeat genContour and mitoMesh for daughter mitochondria after fission. Track the leading-edge retracting from the constriction site, corrected for the motion of the whole mitochondrion, to estimate membrane tension, using a viscoelastic model (16). For membrane tensions estimated from iSIM data, the retraction was fit to an exponential decay function (y=a*exp(bx)), and extrapolated to zero.

To compare the snake and FWHM contours we simulated SIM images using a third-party SIM image generator (17). Different shapes representing mitochondrial constriction sites at an SNR < 2, the typical value at mitochondrial constriction sites, were simulated and used to generate snake and FWHM contours. We observed that although the measured diameters were comparable between the two contours, the snake contour was better at detecting high envelope curvatures (SFig. 6). Hence the analysis was performed on the snake contour. All plots in this work were generated using a third-party function for MATLAB to generate shaded areas representing standard error (18).

To quantify morphological features of mitochondria and after Nocodazole treatment, SIM time-lapse images before and after Nocodazole treatment were analyzed as follows: Mitochondrial membrane potential was analyzed by measuring mean fluorescence intensity of single mitochondria (ROI defined by using Otsu thresholding and Analyse Particles Plugin) on SIM images and subtracting the cytosolic background. The diameter of mitochondria was determined by FWHM measurement profile across individual mitochondria on time-lapse SIM recordings.

ER-mitochondrial overlap was calculated by binarizing both images using the Otsu method in ImageJ. The overlap of the two binarized images was calculated using the Image Calculator AND operation. The overlap was then normalized to the total mitochondrial area.

**Estimating bending energy.**

To determine the area over which to measure the bending energy, we searched for the length scale that maximized the local bending energy density. This was done by considering areas composed of increased numbers of segments around the constriction site (SFig. 2d). The appropriate length scale was then selected by finding the area at which the bending energy density (ratio of bending energy and area) was maximal (SFig. 2e). The bending energy was then considered at this length scale. The length scales did not differ significantly between fissions and reversals (SFig. 2f).

The following equation was then integrated over the area corresponding to the appropriate length scale, to find the bending energy (19) $E_B$:

$$E_B = \frac{\kappa_B}{2} \int J^2 dA$$

Where $\kappa_B$ is the bending rigidity, J the sum of local principal curvatures along the constriction site and A the surface area of the constriction site. The bending rigidity taken to be ~20 kT, as previously reported for lipid bilayers (20). Since we consider a double-membrane system, we take the value of $\kappa_B$= 40 kT.

The above equation was applied to a discretized contour of mitochondria. The contour was subdivided into small segments sampled every ~15 nm. This slightly oversamples the changes in contour, but ensures sufficient sampling at highly bent structures, including constriction sites. The tube curvature was determined by calculating the average radius of the segment. The envelope curvature was taken as the weighted average of the two envelope curvatures, located at each side of the segment. The envelope curvatures were weighted by the length of each side of the segment, and assumed to vary linearly across the two sides of the segment. Finally, to calculate the surface area of the segment, we approximated the shape of the segment as a truncated cone. Therefore, for a single segment, the bending energy was given by:

$$E_B = \frac{\kappa_B}{2} \cdot \left[ \frac{1}{< R_t >_{seg}} + \frac{1}{l_1 + l_2} \left( \frac{l_1}{< R_{e,1} >_{seg}} + \frac{l_2}{< R_{e,2} >_{seg}} \right) \right]^2 \cdot \left[ \pi \left( R_{t,1} + R_{t,2} \right) \sqrt{\left( R_{t,1} - R_{t,2} \right)^2 + h^2} \right]$$

Where $< R_t >_{seg}$ is the average tube radius across the segment, $l_1$ and $l_2$ the side lengths of the two sides of the constriction site, $< R_{e,1} >_{seg}$ and $< R_{e,2} >_{seg}$ the average envelope curvatures of the two sides of the constriction site, $R_{t,1}$ and $R_{t,1}$ the edge tube radii of the segment and $h$ the average length through the center of the segment. This is illustrated in SFig. 2e-f.

**Estimating membrane tension from mitochondrial tubules.**
To estimate mitochondrial membrane tension, we used mitochondrial tubules – short-lived structures pulled out of mitochondria by motor proteins (21, 22) as well as long, thin constriction sites that strongly resembled a membrane pulling experiment. By drawing analogy to membrane pulling experiments (23, 24), the diameter of the tubule d is a readout for membrane tension τ:

$$d = \sqrt{\frac{2\kappa_B}{\tau}} \quad \Rightarrow \quad \tau = \frac{2\kappa_B}{d^2}$$

Where $\kappa_B$ is the bending rigidity, taken as 20 kT for a single membrane (20). The radius of the tube was measured only when the motion of the tube stabilized, and before retraction. Mitochondrial tubules that did not retract and formed a branch on the mitochondrion were not considered. By measuring the diameters of tubules pulled out of mitochondria that subsequently divided, we found d≅200 nm, giving a value of $\tau \cong 4.10^{-6}$ N/m, which provides an independent estimate of membrane tension. The average of the measured tubule diameter was 202 ± 8 nm (mean ± SEM, N=14), above the resolution limit.

**Estimating membrane tension estimate based on retraction speed.** We used a visco-elastic model of the retraction of the mitochondrial membrane after fission to estimate the local membrane tension. The same model was used previously to estimate the membrane tension in the intercellular bridge during cytokinesis (16).

$$\sigma(t) = \int_{-\infty}^{t} \frac{du}{dt}(t = t^*) \times E(t - t^*) dt^* \approx \int_{0}^{t_0} \frac{du}{dt}(t = t_0) \times E(0) dt \approx \frac{v(t_0)}{l_0} \times \eta$$

Where σ is the stress, u the strain, E the relaxation modulus. E(0) is approximated as the effective viscosity η at the moment of fission. $v(t_0)$ is the retraction speed at the moment of fission and $l_0$ the deformation length. $l_0$ was taken as 250 ± 15 nm (mean ± SEM, N=78 recoiling mitochondria from 39 fission events), averaged over all observed fissions. The deformation length was measured by comparing the shape of mitochondria before and after fission. The viscosity was taken as 120 cP. From there, we find the force F by multiplying the stress by the cross-sectional area A:

$$F = \sigma A \cong \sigma \pi r^2$$

Where r is the constriction radius. The membrane tension τ is then estimated by dividing the pulling force by the circumference at the constriction site:

$$\tau = \frac{F}{2\pi r}$$

We suspected that the discrepancy between Methods 1 and 2 was due to suboptimal temporal resolution, which would average out the retraction speed on which retraction-based visco-elastic model is based, decreasing its apparent value. To test this hypothesis, we performed fast live-cell imaging of mitochondria and Drp1, 20x faster than our initial measurements based on the Nikon SIM, using an instant structured illumination microscope (iSIM). We compared the membrane tensions calculated using Method 1 at three different temporal resolutions (SFig. 8c), showing that worse temporal resolution decreased the calculated tension. Using iSIM data with an order of magnitude improvement on temporal resolution, we extrapolated the retraction velocity v at the time of fission by fitting the velocity decay to an exponential function as a function of time t: v=a*exp(bt), where a and b were determined experimentally for each fission event. We repeated the retraction-based measurements, measuring $2.12\pm 1.49 \cdot 10^{-6}$ N/m (mean ± SEM, N=14), comparable to the tubule-based estimate of membrane tension. Thus, the retraction-based estimate of tension quoted in the main text has been independently verified.
Furthermore, the estimate based on tube diameters is likely to be higher since it could only be applied to mitochondria under strong pulling forces with visible tubes, and hence only samples mitochondria with high membrane tension. While the values reported in the manuscript are lower due to low temporal resolution, they still present a real relative difference in membrane tensions of fission events between control and nocodazole treated cells.

**Tension activated model.**
In a model proposed for dynamin-mediated endocytosis, active constriction through GTPase activity increases the membrane elastic energy and brings membranes closer to the energy barrier to fission $E_f$ (26). Given a membrane bending energy $E \in [0, E_f]$, thermal fluctuations allow membranes to stochastically overcome the residual energy barrier $\Delta E$ to fission, with probability $p(E)$ for energy barrier ,

$$\underline{p(E) = exp\left(-\frac{\Delta E}{\lambda}\right) = exp\left(\frac{E - E_f}{\lambda}\right)}$$

Where in the case of thermal fluctuations, $\lambda = k_B T$. The higher the energy of the constricted state, the closer it is to the energy barrier and therefore more likely to divide. Fitting the measured experimental probability of fission (Figure 4A) to the equation above, allowed us to estimate the fluctuation energy $\lambda \sim 90\ k_B T$ and the energy barrier to fission $E_f \sim 250 - 300\ k_B T$.
Fitting equation (2) to both datasets indeed shows that while the barrier to fission remains mostly unchanged, the size of the energy fluctuations was decreased 6-fold (Figure 4B).

**Estimating fission time.**
To estimate the time of fission, we examined the evolution of the constriction site diameter and local Drp1 intensity (SFig. 4a). The fission time was then taken as the offset of the final increase of Drp1 and decrease in constriction diameter, which should correspond to the constriction time before fission.

**Supplementary note 1: Effect of resolution on shape and bending energy**
One of the main features that distinguishes fission from reversal events is the minimal envelope curvature at the constriction site. The resolution limit of ~100 nm for SIM restricts the minimal observable envelope radii to ~50 nm and therefore the maximal observable curvature to $-20*10^{-3}$ $nm^{-1}$. Nevertheless, envelope curvatures for both fissions and reversals remain above the resolution limit of curvature, allowing us to distinguish real differences (Fig. 3d).
The measured constriction diameters remain above the resolution limit of ~100 nm, until a few seconds before maximal constriction (Fig. 3c). We report that in this period during which

constriction sizes are above the resolution limit, constriction diameters are indistinguishable between fissions and reversals (Fig. 3c), while a difference in envelope curvatures can already be observed ~30 s leading up to maximal constriction (Fig. 3d). Furthermore, our STORM measurements, which provide resolution < 100 nm, support that fissions and reversals are not distinguishable by minimum diameter down to ~80 nm (SFig. 2a-c). Therefore, while both fissions and reversals reach the limit of our microscopy techniques, they arrive to this limit in an indistinguishable manner. At their maximal constriction, both fissions and reversals likely involve diameters below the resolution limit of live-cell nanoscopy.

Bending energy is calculated as a function of both envelope and tube curvature – related to the diameter. Therefore, although the calculated bending energies could be underestimated by resolution-limited measurements of the mitochondrial diameter, we are still able to distinguish differences and estimate the energy barrier to fission which is in agreement with theoretical estimates (25, 26) and we expect represents a lower limit on the energy barrier.

**Supplementary note 2: Estimating the energetic contribution of membrane tension**

The contribution of stretching energy was found from two independent estimates described below:

**1) Estimating stretching energy associated with measured membrane tensions?".** Based on mitochondrial geometry and measured membrane tension values, we calculate the stretching energy. Membrane tension can cause small changes in the surface area of the mitochondrial membrane, adding stretching energy to the elastic energy of the membrane. We perform a calculation of the stretching energy based on the measured membrane tension values from iSIM data and mitochondrial tubules (~$4.10^{-6}$N/m). Stretching energy $E_\tau$ due to membrane tension τ and change in surface area ΔA, can be estimated as:

$$E_\tau = \tau \Delta A$$

By modelling a mitochondrion as a cylinder with diameter 1 µm and length 5 µm, and assuming a 0.3% stretch factor (19, 27) (SFig. 4b), we estimated ΔA≅0.05 $\mu m^2$. Combining this estimate with the membrane tension estimate (~$4.10^{-6}$N/m), we obtain that the average energy contribution is ~50 kT.

**2) Experimental estimation of stretching energy**

When tension is applied to a membrane, it causes a change in area resulting in a stretching energy $E_\tau$. Since stretching energy $E_\tau$ scales linearly with membrane tension τ, we have:

$$E_\tau \propto \tau \Rightarrow E_\tau = \alpha\tau \;\Rightarrow \alpha = \frac{dE_\tau}{d\tau} \cong \frac{\Delta E_\tau}{\Delta \tau}$$

We can therefore approximate the constant of proportionality α by looking at the change in stretching energy produced by a change in membrane tension. Specifically, we compared the difference in mean bending energies and mean membrane tensions between control and nocodazole treated cells (SFig. 4c,d). What differentiated these experiments was the loss of mitochondrial membrane tension as described in the main text (Fig. 5). Taken as a consequence of the loss of tension, the difference in bending energies between these two conditions should directly reflect the difference in stretching energies. We estimated the energy due to membrane tension $E_\tau$ as:

$$\boldsymbol{E_\tau} = \left|\frac{\boldsymbol{\Delta E}}{\boldsymbol{\Delta\tau}}\right| \boldsymbol{\tau_{ctrl}} = \left|\frac{\boldsymbol{E_{noco}} - \boldsymbol{E_{ctrl}}}{\boldsymbol{\tau_{noco}} - \boldsymbol{\tau_{ctrl}}}\right| \boldsymbol{\tau_{ctrl}}$$

Where $E_{noco}$, $E_{ctrl}$ are bending energies for nocodazole-treated and control cells respectively, $\tau_{noco}$,$\tau_{ctrl}$ the membrane tensions for nocodazole-treated and control cells respectively. This gives a value of $\boldsymbol{E_\tau}$ of 40 kT, consistent with estimate 1 above.

**Supplementary note 3: Discussion on fluctuation transduction between outer and inner mitochondrial membranes**

From our model, the fission probability is set by two factors: the distance from the energy barrier set by the bending energy and the size of the available fluctuations arising from membrane tension. Let us first consider each factor separately.

Mitochondria are organelles enveloped by inner and outer membranes. If we consider the energetic implications of this architecture, the constrained geometry of the inner mitochondrial membrane implies it should always have the higher bending energies at the constriction site, given the strictly smaller tube radius and therefore higher tube curvature. By assuming a constant distance d between the two mitochondrial membranes (d≈22 nm, (28)), we can relate the local principal curvatures of the outer mitochondrial membrane to the inner as follows (i denotes the inner mitochondrial membrane and o the outer mitochondrial membrane) (SFig. 5a):

$$R_{e,i} = R_{e,t} + d$$

$$R_{t,i} = R_{t,o} - d$$

Where $R_e$ and $R_t$ are the envelope and tube radii of curvature respectively. This is supported by the observation from electron microscopy that cristae are excluded from constriction site (29). From this we can compute a readout for the bending energy density at a single point along the membrane (neglecting the bending rigidity κ) (SFig. 5b):

$$\epsilon_b = \left(\frac{1}{R_e} + \frac{1}{R_t}\right)^2$$

The consequence is that for most geometries observed from our data (Fig 1,3), the inner mitochondrial membrane will intrinsically have a higher bending energy and will therefore be closer to the energy barrier to fission (smaller residual energy ΔE). By comparing the bending energies of the two membranes, we can see that on average the inner membrane will have almost a two-fold higher bending energy (SFig. 5c), and be closer to the energy barrier to fission.

What effect would differences in membrane tension fluctuations have on the fission probability of the inner and outer membranes? Any membrane tension-inducing force arising from the surroundings acting on the mitochondria would be exerted through the outer mitochondrial membrane. We expect that coupling, which could arise from proteins spanning the two membranes such as Tim23 (30), would transduce a part of the force to the inner membrane.

The probability of fission P is related to the residual energy (distance to the energy barrier) ΔE, and the inverse of the fission time $t_f$ by:

$$P \propto \frac{1}{t_f} \propto \exp\left(-\frac{\Delta E}{\lambda}\right)$$

Where λ reflects the magnitude of the fluctuations. We introduce a coupling variable $\omega \in [0,1]$ such that $\lambda_i = \omega\lambda_o$ where $\lambda_i$ reflects the fluctuations transduced to the inner membrane from the outer membrane ($\lambda_o$). In this case, we can express the probabilities as

$$P_o \propto \frac{1}{t_{f,o}} \propto \exp\left(-\frac{\Delta E}{\lambda_o}\right)$$

$$P_i \propto \frac{1}{t_{f,i}} \propto \exp\left(-\frac{\Delta E}{\lambda_i}\right) = \exp\left(-\frac{\Delta E}{\omega\lambda_o}\right)$$

We can then express $P_i$ as a function of $P_o$ and ω, where the constant of proportionality is 1:

$$P_i = P_o^{1/\omega}$$

In this case, unless the coupling is perfect (ω=1), the inner mitochondrial membrane will always be less likely to undergo fission than the outer (SFig. 5d). The inner mitochondrial membrane would therefore need to achieve higher energies to overcome the energy barrier, given the lower scale of available fluctuation energies.

We wondered whether the higher bending energy of the inner membrane could compensate for the imperfect force transduction, so let us now consider the two factors together. We can compute again the probability of fission for the two mitochondrial membranes:

$$P_o \propto \frac{1}{t_{f,o}} \propto \exp\left(-\frac{\Delta E}{\lambda_o}\right)$$

$$P_i \propto \frac{1}{t_{f,i}} \propto \exp\left(-\frac{\Delta E}{\lambda_i}\right) = \exp\left(-\frac{\Delta E}{\omega\mu\lambda_o}\right)$$

Where μ represents the ratio of residual energies. This implies that the inner mitochondrial membrane can still have a higher probability of fission, even if the coupling to the outer membrane is not perfect (SFig. 5e,f) and therefore will be more likely to undergo fission (SFig. 5e,f). For factors of $\mu>1$, there will exist cases of imperfect coupling ($\omega<1$), which will still give the inner and outer membranes equal fission probabilities (SFig. 5g-i). Furthermore, for a fixed ratio of bending energies μ we can identify 3 regimes: (I) $P_i<P_o$ below a critical coupling ($\omega<\omega_c$), (II) $P_i>P_o$ for ($\omega>\omega_c$), and (III) $P_i=P_o$ at critical coupling ($\omega=\omega_c$).

The necessary coupling for a combination of $P_i$, $P_o$, μ can be found as:

$$\omega = \frac{\log P_o}{\mu \log P_i}$$

Therefore, as the inner membrane gets relatively closer to the energy barrier (increasing μ), the required coupling will decrease (SFig. 5j-l). Critical coupling, $\omega_c$, corresponding to $P_i=P_o$ is given by:

$$\omega_c = \frac{1}{\mu}$$

Therefore, even in cases where the coupling is not perfect ($\omega<1$), our model predicts that the inner membrane can still undergo fission first. Furthermore, in reality the ratio of bending energies μ is expected to be even higher than 2, suggesting that the regime where the inner membrane undergoes fission first can exist at even much lower degrees of coupling.

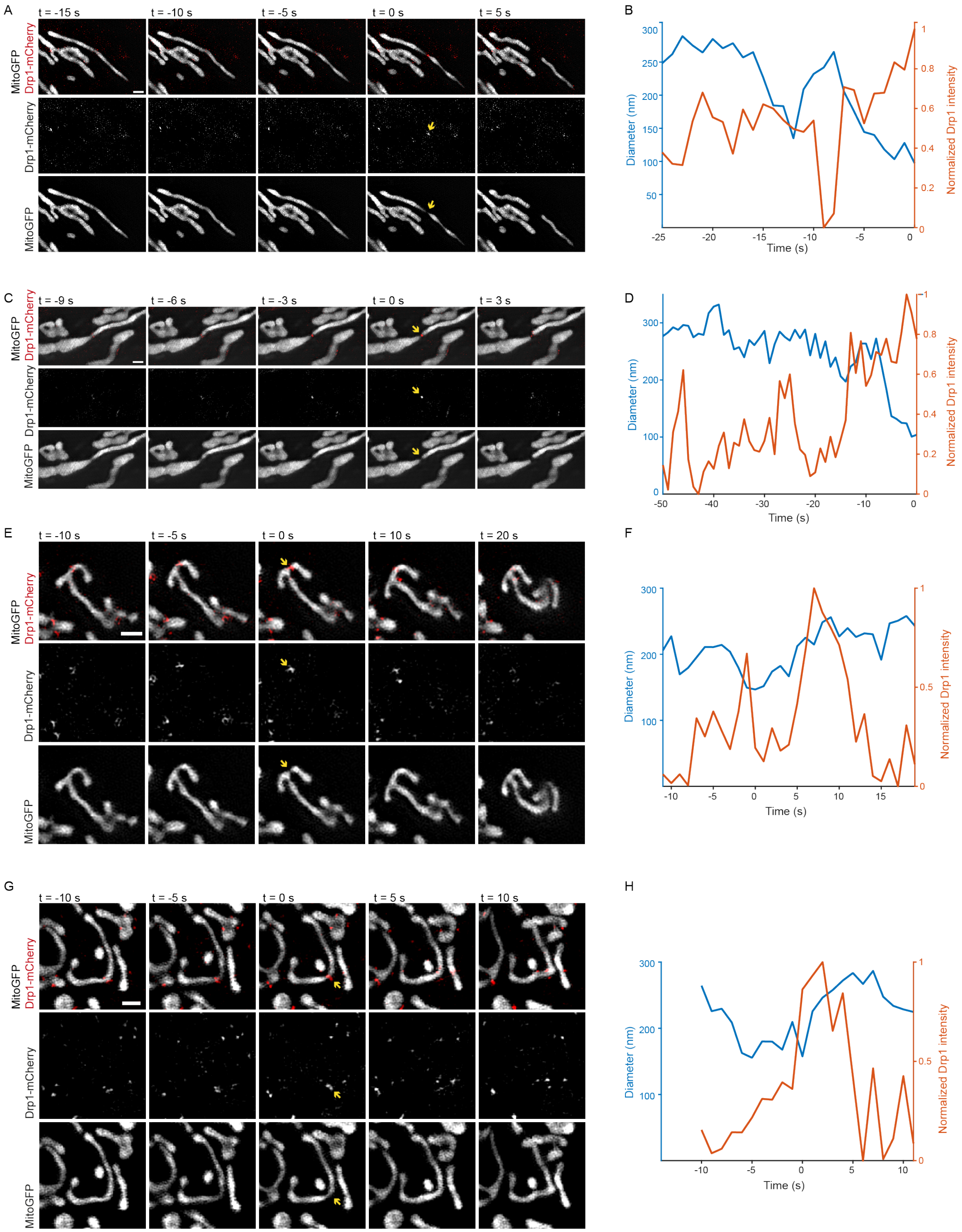


**Fig. S1. Examples of fission and reversal events.** (A,C) Time-lapse SIM imaging of COS-7 cells transiently transfected with mito-GFP (grey) and mCh-Drp1 (red) showing exemplar fission events. (B,D) Corresponding measurements of the constriction diameter dynamics (blue, left axis) and evolution of the normalized integrated intensity of Drp1 at the constriction site (red, right axis) of fission events. (E,G) Time-lapse SIM imaging of COS-7 cells transiently transfected with mito-GFP (grey) and mCh-Drp1 (red) showing exemplar reversal events. (F,H) Corresponding measurements of the constriction diameter dynamics (blue, left axis) and evolution of the normalized integrated intensity of Drp1 at the constriction site (red, right axis) of reversal events. Images were adjusted for contrast. Scale bars represent 1 µm in each panel.

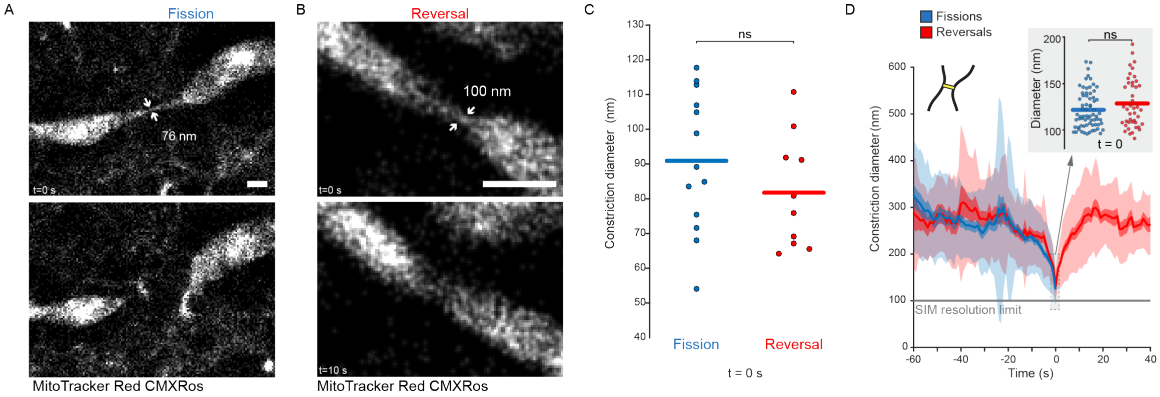


**Fig. S2. Live-cell STORM imaging of fate of mitochondrial constriction sites and energetics of the constriction site.** (A,B) Time-lapse live-cell STORM imaging of COS-7 cells labelled with MitoTracker Red CMXRos showing a (A) fission and a (B) reversal event. Arrows indicate measured diameter values. (C) Distribution of minimum constriction diameters for fissions (N=13) and reversals (N=10) measured with STORM. Horizontal line represents the mean. (D) Binned curves showing the evolution of the constriction diameter for fissions (blue, n=69) and reversals (red, n=43). Inset shows the distribution of minimal diameters of fission and reversals at t=0 (horizontal line shows mean).

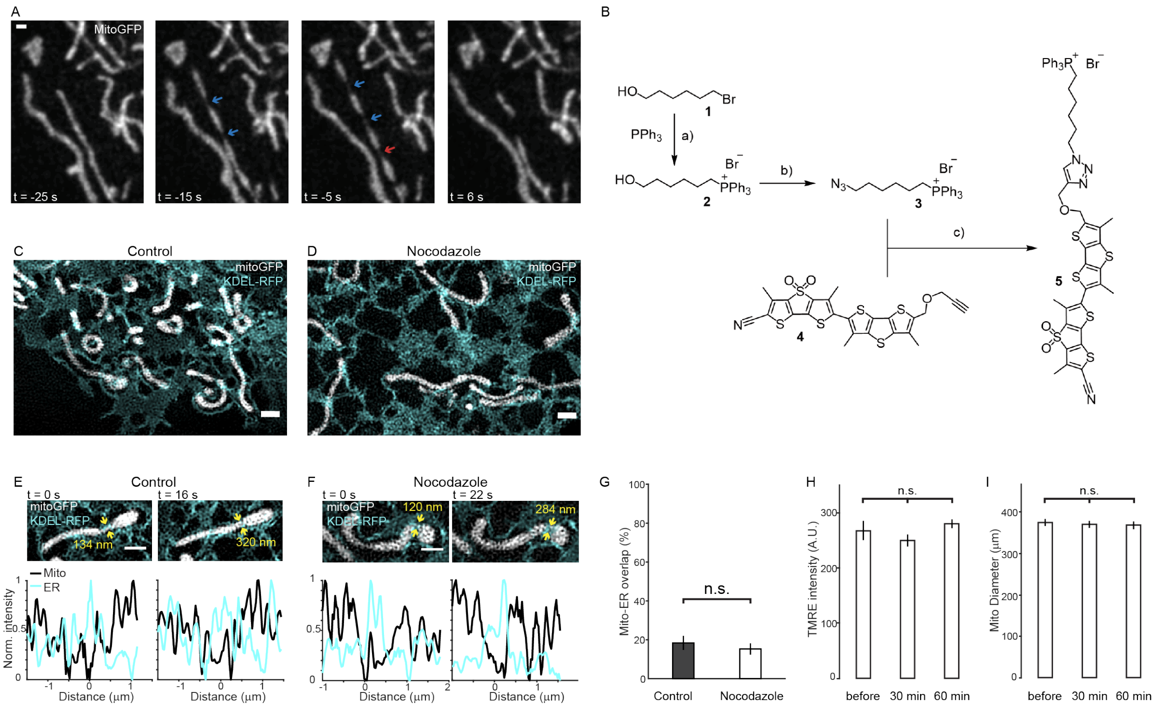


**Fig. S3. FliptR synthesis and nocodazole controls.**

(A) Example of pearling modes observed on mitochondria. Blue arrows indicate constriction sites that result in fission and red arrows indicate the ones that reverse. Scale bar: 500 nm. (B) Toluene, reflux, 6 h, 30%. (b) 1. MsCl, $NEt_3$, THF, 0 °C to rt, 30 min; 2. $NaN_3$, THF/DMF, 60 °C, 2 h, 50%. (c) $CuSO_4.5H_2O$, Na-ascorbate, TBTA, $CH_2Cl_2$, DMSO, $H_2O$, rt, 2 h, 59%. Compounds 2,3 and 5 were synthesized and purified according to procedures that will be reported elsewhere in another manuscript S1 in due time. Compound 4 was synthesized and purified according to procedures described in ref. (31). (C,D) Representative images showing the organization of the ER between control (C) and Nocodazole treatment (D) in COS-7 cells transfected with Mito-GFP (grey) and KDEL-RFP (cyan). (E,F) Time-lapse live-cell SIM imaging of mitochondria and ER in COS-7 cells transfected with Mito-GFP and KDEL-RFP showing reversible constrictions occurring in the presence of the ER in both (E) control and (F) nocodazole treated cells. Images were adjusted for contrast. (G) Quantification of mitochondrial-ER overlap as a percentage of the mitochondrial surface. Lack of statistical significance between control (N=9) and Nocodazole (N=10) calculated by computing the 95% confidence interval for the difference between means. (H) Quantitative analysis of TMRE staining before and 30 min/60 min after Nocodazole treatment confirms no significant difference of mitochondrial membrane potential after drug treatment ($N \geq 250$ mitochondria for each group, error bars represent ±SEM). Statistical significance calculated by Kruskal-Wallis ANOVA (I) Quantification of mitochondrial diameter before and 30 min/60 min after Nocodazole treatment confirms no change in mitochondrial diameter and absence of mitochondrial swelling (N=200 mitochondria for each group, error bars represent ±SEM). Statistical significance calculated by Kruskal-Wallis ANOVA. Scale bar represents 1 μm in all panels. n.s. indicated $p \geq 0.05$.

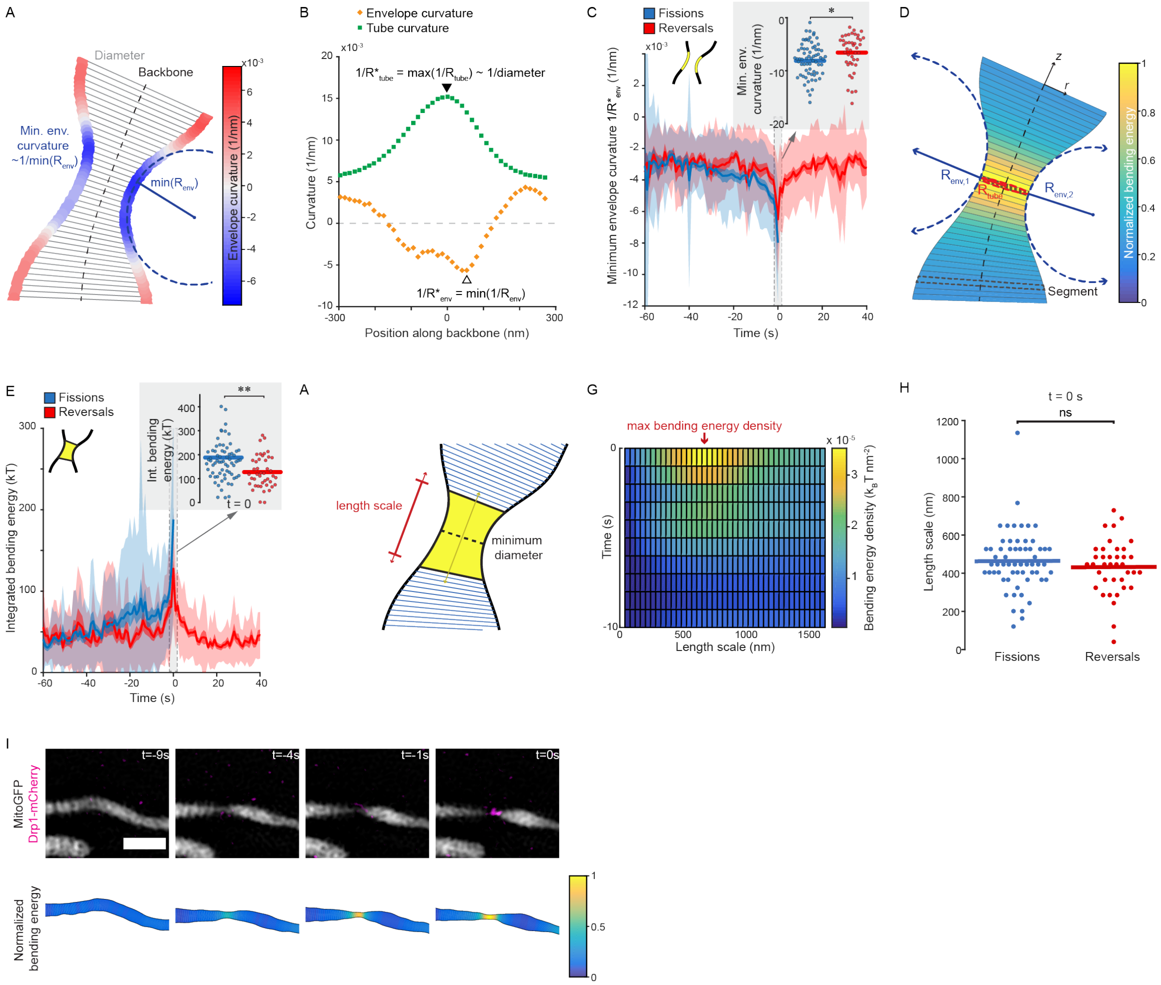


**Fig. S4. Curvature and bending energy at the constriction site for fissions and reversals.** (A) Schematic showing the local envelope curvature at a constriction site (colormap). (B) Tube curvature and envelope curvature measured along the backbone at the constriction site in (a). (C) Binned curves showing the evolution of envelope curvature at the constriction site for fissions (blue, n=69) and reversals (red, n=43). Shaded regions represent SD (light shade) and SEM (dark shade). Inset shows the distribution of minimal envelope curvatures of fission and reversals at t=0 (horizontal line shows mean). (D) Schematic showing the bending energy measurement. The local envelope curvature R_env was computed as the local mean of opposing sides R_(env,1) and R_(env,2). Analysis of the local tube (R_tube) and envelope (R_env) radii for individual segments (striped region) allowed estimation of the local bending energy (colormap). (E) Binned curves for the calculated local bending energies over time for fissions (blue, n=70) and reversals (red, n=43). Shaded regions represent SD (light shade) and SEM (dark shade). Inset shows the distribution of integrated bending energies at the constriction site of fission and reversals at t=0 (horizontal line shows mean). (F) Cartoon illustrating how the length scale was determined by considering increasing areas around the constriction site. (G) The length scale was chosen by maximizing the local bending energy density. (H) Distributions of measured length scales for fissions (N=61) and reversals (N=38). Statistical significance calculated by a paired t-test: n.s. indicated p≥0.05. (I) Timelapse showing the raw data (top) and estimated local bending energies (bottom, colormap) of a Drp1 mediated constriction site resulting in fission. Scale bar: 1um. Statistical significance calculated by 2-tailed Mann Whitney U test: n.s. P>0.05, *P<0.05, **P<0.01.

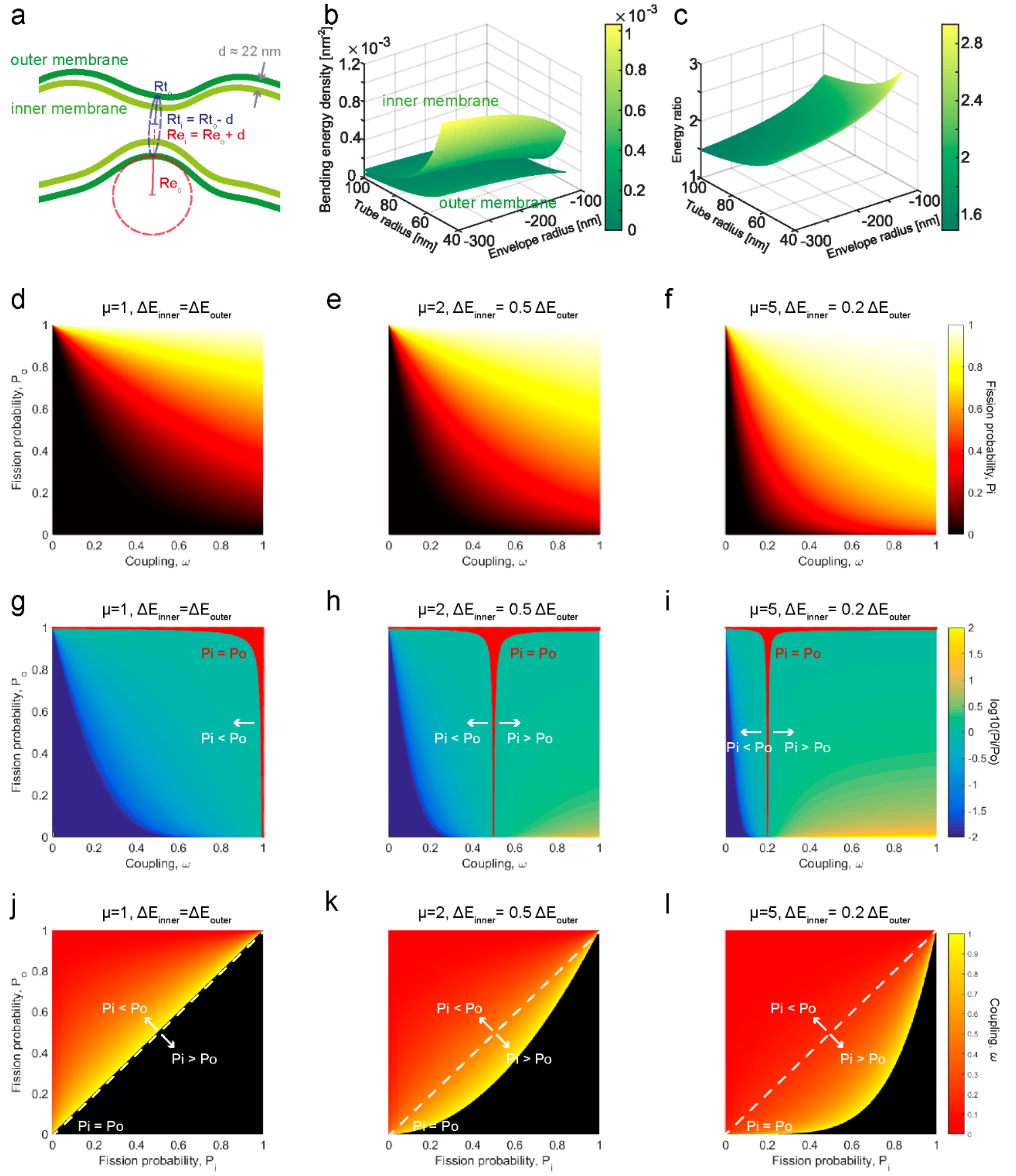


**Fig. S5. Model for inner and outer mitochondrial membranes.** (a) Schematic representation showing the geometries of the outer and inner mitochondrial membranes, as a function of the local tube radius ($R_{t,o}$), envelope radius ($R_{e,o}$) of the outer membrane and distance d between the two membranes. (b) 3D surface plot showing the bending energy densities for the inner and outer mitochondrial membranes, as a function of the outer membrane radii of curvature. (c) 3D surface plot showing the ratio between the bending energies of the inner to outer mitochondrial membranes, as a function of the outer membrane radii of curvature. (d-f) Fission probability for the inner mitochondrial membrane ($P_i$, colormap) as a function of the fission probability for the outer mitochondrial membrane ($P_o$) and the degree of coupling ω between the two. The inner membrane is considered (d) at the same distance ($\mu$=1), (e) twice closer ($\mu$=2) and (f) five times closer ($\mu$=5) to the energy barrier than the outer membrane. (g-i) Log of the ratio of probabilities of fission for the inner membrane to outer mitochondrial membrane ($\log(P_i/P_o)$, colormap) as function of the fission probability for the outer mitochondrial membrane and degree of coupling $\omega$. The inner membrane is considered (g) at the same distance ($\mu$=1), (h) twice closer ($\mu$=2) and (i) five times closer ($\mu$=5) to the energy barrier than the outer membrane. Red marks the region where the inner and outer mitochondrial membranes have similar probabilities of fission (within 0.5%). (j-l) Optimal coupling values ($\omega$, colormap), as a function of the inner and outer membrane fission probabilities. The inner membrane is considered (j) at the same distance ($\mu$=1), (k) twice closer ($\mu$=2) and (l) five times closer ($\mu$=5) to the energy barrier than the outer membrane. White dashed line marks $P_i$=$P_o$. Black indicates that no coupling value can give the corresponding inner and outer membrane fission probabilities.

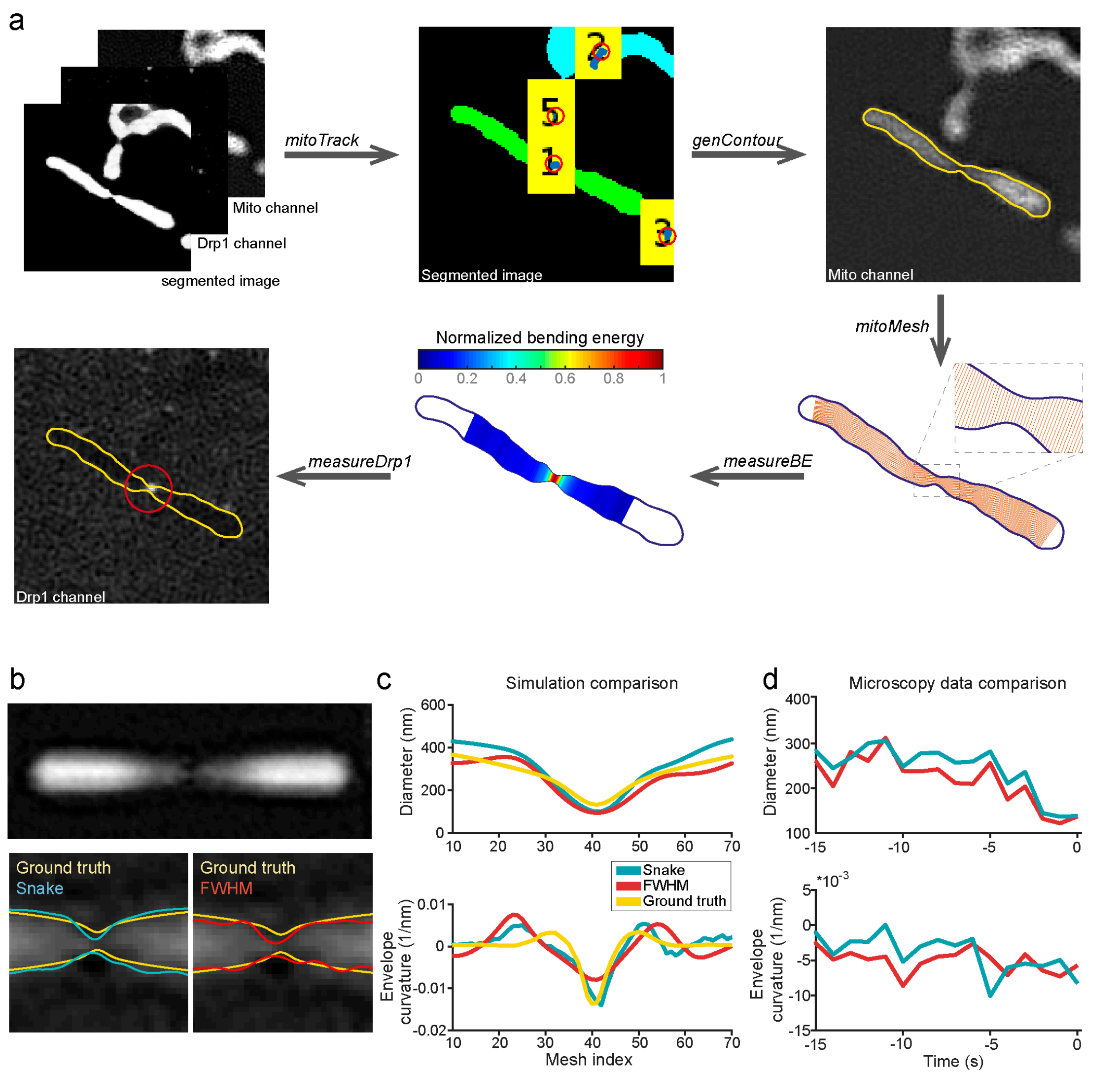


**Fig. S6. MitoWorks analysis.** (a) Schematic representation of the MitoWorks analysis workflow. Two raw channels and a segmented image are chosen as inputs. The mitochondria were then tracked, and a mitochondrion of interest chosen by the user. This mitochondrion is then contoured, from which MitoWorks creates a mesh and a skeleton. This allows the shape and energetics of the constriction to be analyzed over time. Finally, the Drp1 signal at the constriction site is measured. (b) Simulated SIM image of a constricted mitochondrion was created based on a ground truth (yellow outline) to test the performance of the snake (blue outline) and FWHM (red outline) contours. (c) Measurement of constriction diameters (top) and envelope curvatures (bottom) for Snake and FWHM contours, compared to the ground truth structure (yellow line). (d) Comparison of snake and FWHM contour from real microscopy data.

**Movie S1.** Example fission event. Time-lapse SIM imaging of COS-7 cells transiently transfected with mito-GFP (grey) and mCh-Drp1 (red) showing a fission event. Time between frames: 1s, scale bar: 1 μm.

**Movie S2.** Example reversal event. Time-lapse SIM imaging of COS-7 cells transiently transfected with mito-GFP (grey) and mCh-Drp1 (red) showing a reversal event. Time between frames: 1s, scale bar: 1 μm.

**Movie S3.** Time-lapse showing the evolution of the bending energy during a Drp1 mediated mitochondrial constriction. Time between frames: 1s, scale bar: 1 μm.

**Movie S4.** Example recoil after fission. Time-lapse SIM imaging of COS-7 cells transiently transfected with mito-GFP (grey) and mCh-Drp1 (red) showing a fission event with significant recoil after fission. Time between frames: 1s, scale bar: 1 μm.

**Movie S5.** Example recoil after fission. Time-lapse SIM imaging of COS-7 cells transiently transfected with mito-GFP (grey) and mCh-Drp1 (red) showing a fission event with significant recoil after fission. Time between frames: 1s, scale bar: 1 μm.
